# Membrane-remodeling protein ESCRT-III homologs incarnate the evolution and morphogenesis of multicellular magnetotactic bacteria

**DOI:** 10.1101/2022.11.08.515611

**Authors:** Wenyan Zhang, Jianwei Chen, Jie Dai, Shiwei Zhu, Hugo Le Guenno, Artemis Kosta, Hongmiao Pan, Xin-Xin Qian, Claire-Lise Santini, Nicolas Menguy, Xuegong Li, Yiran Chen, Jia Liu, Kaixuan Cui, Yicong Zhao, Guilin Liu, Eric Durand, Wei-Jia Zhang, Alain Roussel, Tian Xiao, Long-Fei Wu

## Abstract

Endosomal sorting complex required transport (ESCRT) III proteins are essential for membrane remodeling and repair across all domains of life. Eukaryotic ESCRT-III and the cyanobacterial homologs PspA and Vipp1/Imm30 remodel membrane into vesicles, rings, filaments and tubular rods structures. Here our microscopy analysis showed that multicellular bacteria, referred to as magnetoglobules, possess multiple compartments including magnetosome organelles, polyphosphate granules, vesicles, rings, tubular rods, filaments and MVB-like structures. Therefore, membrane remodeling protein PspA might be required for the formation of these compartments, and contribute to the morphogenesis and evolution of multicellularity. To assess these hypotheses, we sequenced nine genomes of magnetoglobules and found a significant genome expansion compared to unicellular magnetotactic bacteria. Moreover, PspA was ubiquitous in magnetoglobules and formed a distinct clade on the tree of eubacterial and archaeal ESCRT-III. The phylogenetic feature suggested the evolution of magnetoglobules from a unicellular ancestor of deltaproteobacterium. Hetero-expression of ellipsoidal magnetoglobule *pspA2* gene alone in *Escherichia coli* resulted in intracellular membrane aggregation. GFP fusion labeling revealed polar location of PspA2 in rod-shaped unicells and regular interval location in filamentous cells. Cryo-electron tomography analysis showed filament bundle, membrane sacculus, vesicles and MVB-like structure in the cells expressing PspA2. Moreover, electron-dense area with a similar distribution as GFP-PspA2 foci in filamentous cells changed the inward orientation of the septum, which might interfere with the cell division. Collectively, these results show the membrane remodeling function of magnetoglobule PspA proteins, which may contribute to morphogenesis and the evolution of multicellularity of magnetotactic bacteria.

## Introduction

During the evolution, eukaryotic cells have developed various internal compartments to efficiently fulfill defined functions crucial for life. One of them, endosomes, is primarily intracellular sorting organelle required for the trafficking of proteins and lipids among the compartments. The endosomal sorting complex required transport (ESCRT) machinery was originally identified for its function in delivering cargo from the plasma membrane or *trans*-Golgi to the vacuole or lysosome via the generation of multivesicular body (MVB) by deforming the endosomal-limiting membrane inward ^1^. MVB is a hallmark of eukaryotic cells and their formation depends on ESCRT machinery including ESCRT-III proteins that function in membrane remodeling, repairing and membrane abscission in cytokinesis in eukaryotic cells ^2^. Recently, phylogenetic and structural studies have shown that bacterial Vipp1/Imm30 and PspA proteins are members of the ESCRT-III membrane-remodeling superfamily, which plays pivotal role in maintaining membrane integrity and thylakoid biogenesis, and is capable of remodeling lipid bilayers in vitro ^3-6^. This finding paves the way toward a better understanding of the mechanism of compartmentalization in the primitive cells of bacteria.

Magnetotactic bacteria (MTB) are a phylogenetically, physiologically and morphologically heterogeneous group of Gram-negative bacteria ^7-9^. They all produce magnetosomes, which are bacterial organelles composed of single domain magnetic nanocrystals enclosed in membrane. Magnetosomes confer a magnetic dipolar moment to cells and allow them aligning in and swimming along magnetic field lines. The formation of magnetosomes is a genetic-controlled and enzyme-catalyzed process. Cryo-electron tomography (CET) analysis shows that magnetosome membrane is either continuous with or derived from the cytoplasm membrane ^10,11^. Magnetosome biogenesis starts with invagination of cytoplasmic membrane through a protein crowding process ^12-14^. Genetic and molecular studies indicate that MamB and MamM are important for magnetosome membrane formation in *Magnetospirillum magnetotacticum* AMB-1 ^14,15^ and *Magnetospirillum gryphiswaldense* MSR-1 ^16-18^. Besides magnetosomes, MTB cells also contain phosphate or lipid granules and ferrosomes ^14,19^. Formation of ferrosomes depends on ferrosome-associated (Fez) proteins that are well conserved in diverse bacteria ^19^. Whether the membrane remodeling proteins ESCRT-III are involved in formation of these MTB organelles and granules has not been analyzed.

MTB exhibit myriad morphologies including cocci-ovoid, rods, vibrios, spirilla and more complex multicellular forms. Spherical magnetotactic multicellular aggregates (MMAs) were first reported by Farina et al. ^20^. Later, Rodger et al. described many-celled magnetotactic prokaryotes (MMP) with similar morphology and swimming behavior as MMAs ^21^. Typically, 15–45 bacterial cells arrange with a helical geometry in a spherical multicellular entity with an internal acellular compartment ^22^. This morphotype of magnetotactic organisms is observed worldwide and referred to as spherical or rosette/mulberry-shaped magnetotactic multicellular prokaryote (sMMP). Genomic analysis of a spherical uncultured magnetotactic prokaryote, *Candidatus* Magnetoglobus multicellularis, revealed several proteins, including hemagglutinin-like proteins, adhesion-like proteins, glycoprotein and integrin- and fibronectin-like proteins. These proteins were proposed to be involved in multicellular morphogenesis and development of multicellular organization in this organism ^23^. Based on the nomenclature of ‘Magnetoglobus’, the multicellular magnetotactic prokaryotes can be referred to, with a shorter name, as magnetoglobules. Genome of another spherical magnetoglobule, *Candidatus* Magnetomorum sp. strain HK-1, has been sequenced ^24^. Interestingly, it possesses two paralogous copies with highest similarity to either greigite-type magnetosome genes from *Candidatus* M. multicellularis or magnetite-type magnetosome genes from unicellular MTB. Besides the spherical MMPs, we have identified another morphotype, the ellipsoidal or pineapple-shaped magnetoglobules (eMMPs) in the Mediterranean Sea, the China Sea and the Pacific Ocean ^25-29^. Approximately 60 phylogenetically identical cells assemble into a one-layer ellipsoidal entity ^30^. Both morphotypes of magnetoglobules have the center acellular compartment or the core lumens containing vesicles probably required for molecule and information exchange among the cells. Magnetoglobules reproduce without individual cell stage through periphery-to-center unilateral invaginations of constituent cell membrane with an unknown mechanism ^30-32^. Magnetoglobules are phylogenetically, morphologically and reproductively distinct from the extensively studied multicellular cyanobacteria, actinobacteria and myxobacteria ^30^. The origin of the magnetoglobules as well as the involvement of ESCRT-III in the formation of the compartments and multicellular morphogenesis of magnetoglobules remain unknown. Here, to better understand the evolution of magnetoglobule multicellularity, we sequenced the genomes of five spherical and four ellipsoidal magnetoglobules, and analyzed them together with the 44 unicellular MTB genomes and the two spherical magnetoglobules genomes available in the Genome Taxonomy Database (GTDB). The phylogenetic tree suggests that magnetoglobules evolved from a unicellular deltaproteobacterial ancestor. Absence of *pspA* from several taxonomic groups of unicellular MTBs indicated that PspA is unlikely required for magnetosome membrane formation. All magnetoglobules possessed PspA that clustered in a detached clade. Heterologous expression of the representative *pspA* genes of ellipsoidal magnetoglobules in *E. coli* resulted in filamentous cell morphology, aggregation of membrane, and formation of vesicles, and multivesicular body-like structure. Collectively, our results show a membrane remodeling function of PspA that contribute to intracellular compartment formation and might be involved in multicellular morphogenesis of magnetoglobules.

## Results

### Magnetoglobules are rich in intracellular granules

Since the discovery of magnetoglobules four decades ago, despite efforts of various laboratories, the organisms have still not been cultivated. The study of magnetoglobules is mainly carried out through imaging, biophysical, taxonomic and comparative genomics analyses. Scanning transmission electron microscopy in combination with annular darkfield imaging (STEM-HAADF) offers a better depth of observation field and a better contrast of the intracellular vacuoles and elements compared to conventional transmission electron transmission (TEM) dark-filed imaging. It is a highly suitable method for imaging the thick magnetoglobules. STEM-HAADF inspection clearly showed magnetosome chains and vacuolar granules in magnetoglobules (Figure 1, A). Elemental map of these intracellular components was performed with energy-dispersive X-ray spectroscopy acquisition in parallel with STEM imaging (STEM-XEDS). Granules containing oxygen, magnesium, potassium, calcium and phosphor were scattered throughout the magnetoglobules (Figure 1, B to I). They might be polyphosphate granules with incorporation of Mg, K, Ca cations as observed in different ectomycorrhizal fungi ^33^.

**Figure 1.**
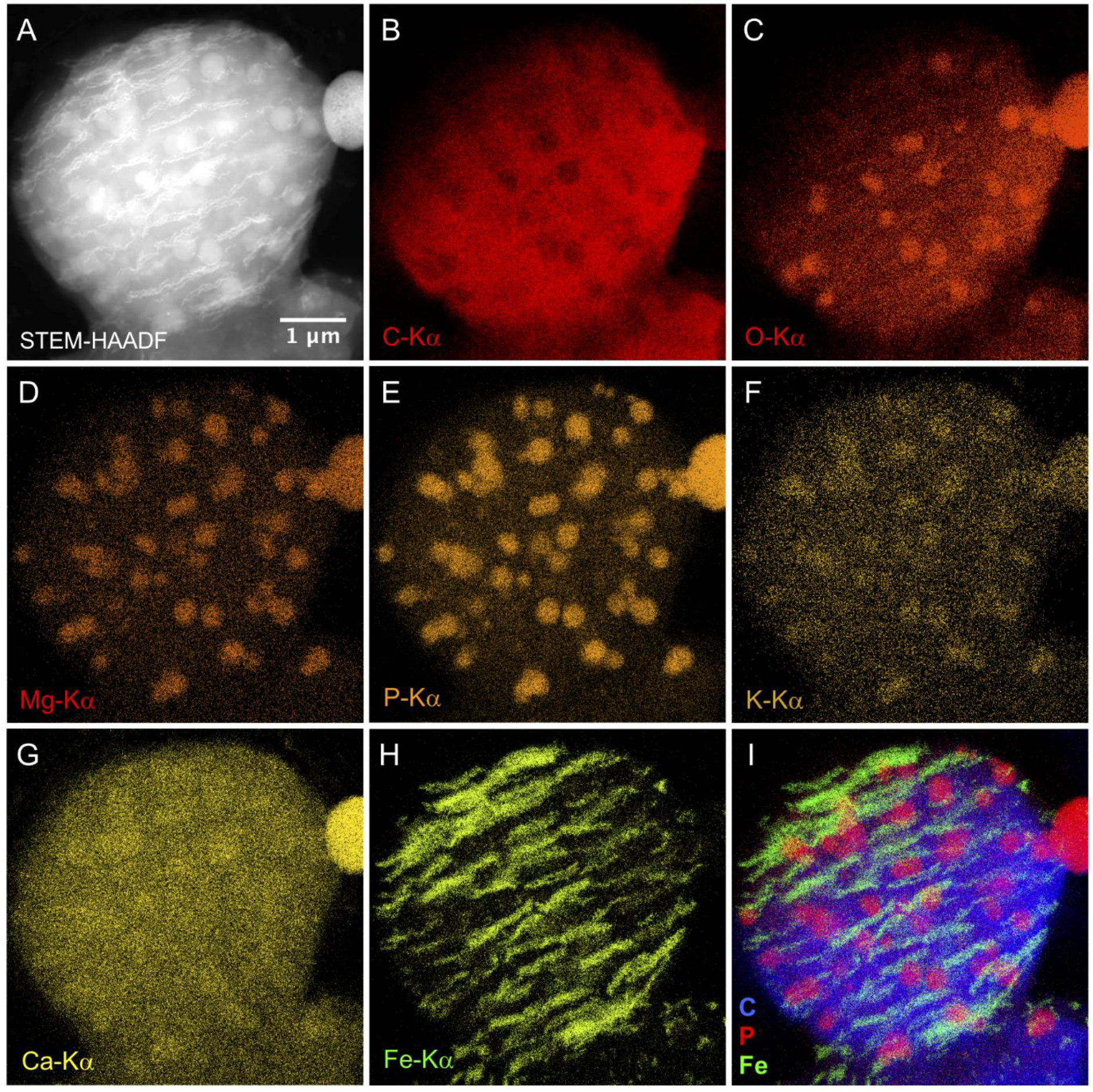
Magnetosome chains and vacuolar granules in ellipsoidal magnetoglobules. STEM-HAADF (A) and STEM-XEDS elemental maps (B – H) of a magnetoglobule. Composite image of Carbon, Phosphorus and Iron (I).

### Magnetoglobules may undergo extensive membrane remodeling process

TEM inspection of ultrathin sections of high-pressure freezing/freeze substitution fixation (HPF/FS) samples showed vesicles with diameters from 35.5 nm to 57.8 nm in core lumen (red arrows in Figure 2A), in the out-surface matrix (Figure 2A and 2C) and in the periplasm of cells where they appeared like multivesicular body (Figure 2D). Nil red-stainable lipid granules were surrounded by a slightly electron-dense circle, which suggested a polymer coating on the granules (Figure 2A). Periphery bars exhibited a filamentous structure with a width of 30.0 ± 9.0 nm and length of 363.4 ± 146.7 nm (yellow arrows in Figure 2A and 2B). Remarkably, the cytoplasm of the cells was filled with “C” and “S” shaped open rings (blue arrows in Figure 2A), or tubular rods with width of 72.8 ± 6.6 nm and length of 337.7 ± 103.4 nm (green arrows), which recalls the morphology of the Vipp1 oligomer observed in the cyanobacterium *Synechocystis* PCC 6803 ^34^. Formation of these structures may depend on an extensive membrane remodeling.

**Figure 2.**
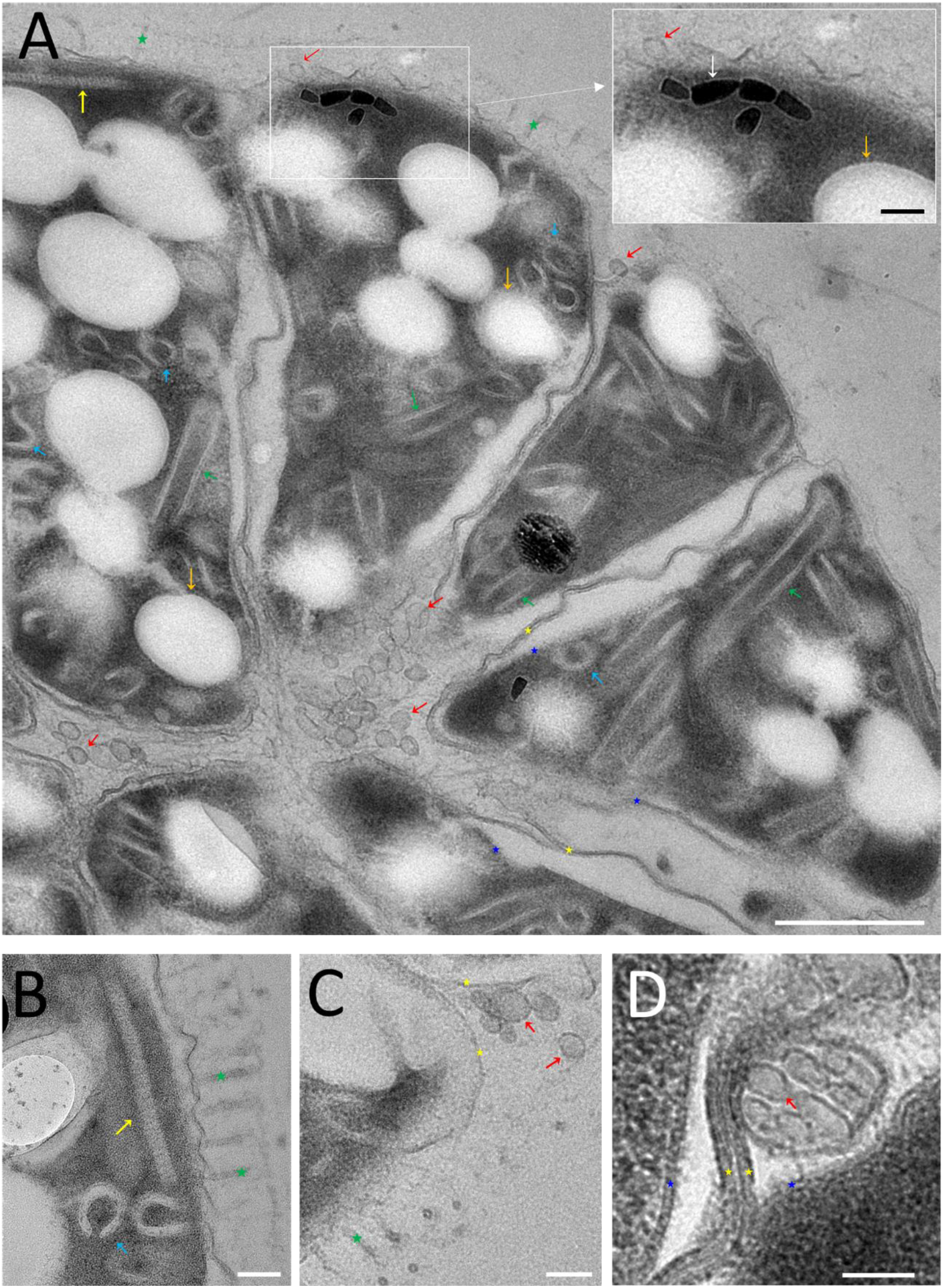
Vesicles and filamentous structures in ellipsoidal magnetoglobules. Micrographs of ultrathin sections of HPF/FS-fixed ellipsoidal magnetoglobules show vesicles (red arrows) in the core lumen (A), in the out-surface matrix (A and C), and appeared as multivesicular body in the periplasm of cells (D). Magnetosomes are indicated by white arrow in (A). Filaments are shown by yellow arrows (A and B). Orange arrows point to Nil red-stainable lipid granules. Intracellular tubular rods are indicated by green arrows (A). Cyan arrows show “C” or “S” rings (A and B). Yellow and blue asterisks show the outer and inner membranes, respectively (A, C and D). Green asterisks show the flagella (A, B and C). Scale bars indicate 500 nm in (A) and 100 nm in (A)-inset and other panels.

### Genome sequence of magnetoglobules

To analyze the presence of membrane-remodeling gene in magnetoglobules, we isolated magnetoglobules from environmental samples collected from the China Sea and the Mediterranean Sea and sequenced genomes of 4 ellipsoidal and 5 spherical magnetoglobules (Table 1). The assembled genomes had an average completeness of 94.9 ± 2.4% and contamination of 3.0 ± 1.2%. The four ellipsoidal magnetoglobules with average genome size 10.1 Mbp (9.8 to 10.8 Mbp) and GC content about 34% were all assembled into pseudo-chromosomal level genome with only one scaffold, and the number of contigs ranged from 216 to 622. Compared to the ellipsoidal magnetoglobule genomes, those of spherical magnetoglobules showed higher variations in size (ranging from 8.8 to 11.4 Mbp) and GC content (29.9 to 36.4%). Based on the 16S rRNA gene identity, genomic average nucleotide identity (ANI) and the GTDB-Tk taxonomic annotation using the relative evolutionary divergence (RED), the spherical magnetoglobules were classified into 5 species of 3 genera and the 4 ellipsoidal magnetoglobules into 3 species of the same genus (Table S1), using the cutoff values (Same species: 16S rRNA gene sequence identity > 97%, ANI > 95%, RED > 0.85, Different genus: 16S rRNA gene sequence identity < 92%, ANI < 83%, 0.7 < RED < 0.85) published in ^35-37^. The higher taxonomic diversity of spherical magnetoglobules was consistent with the bigger difference of their genome sizes compared to the ellipsoidal magnetoglobule genomes. The five spherical magnetoglobules were classified into *Candidatus* Magnetopila (for ZJ64 and ZJ12) and *Candidatus* Magnetoradiorum (ZJW7) two novel genera, and *Candidatus* Magnetomorum zhanjiangroseum (ZJ63) and *Candidatus* Magnetomorum huiquanroseum (QDA1) two novel species (Table 1). The four ellipsoidal magnetoglobules were affiliated to *Candidatus* Magnetananas genus, two previously named species *Candidatus* Magnetananas rongchenensis (RCG1) and *Candidatus* Magnetananas tsingtaoensis (QDG1) and a new species *Candidatus* Magnetananas bruscensis (for SF-25 and SF-35). The five spherical and four ellipsoidal magnetoglobules all belonged to Deltaproteobacteria, Desulfobacterales, Desulfobacteraceae.

**Table 1.**
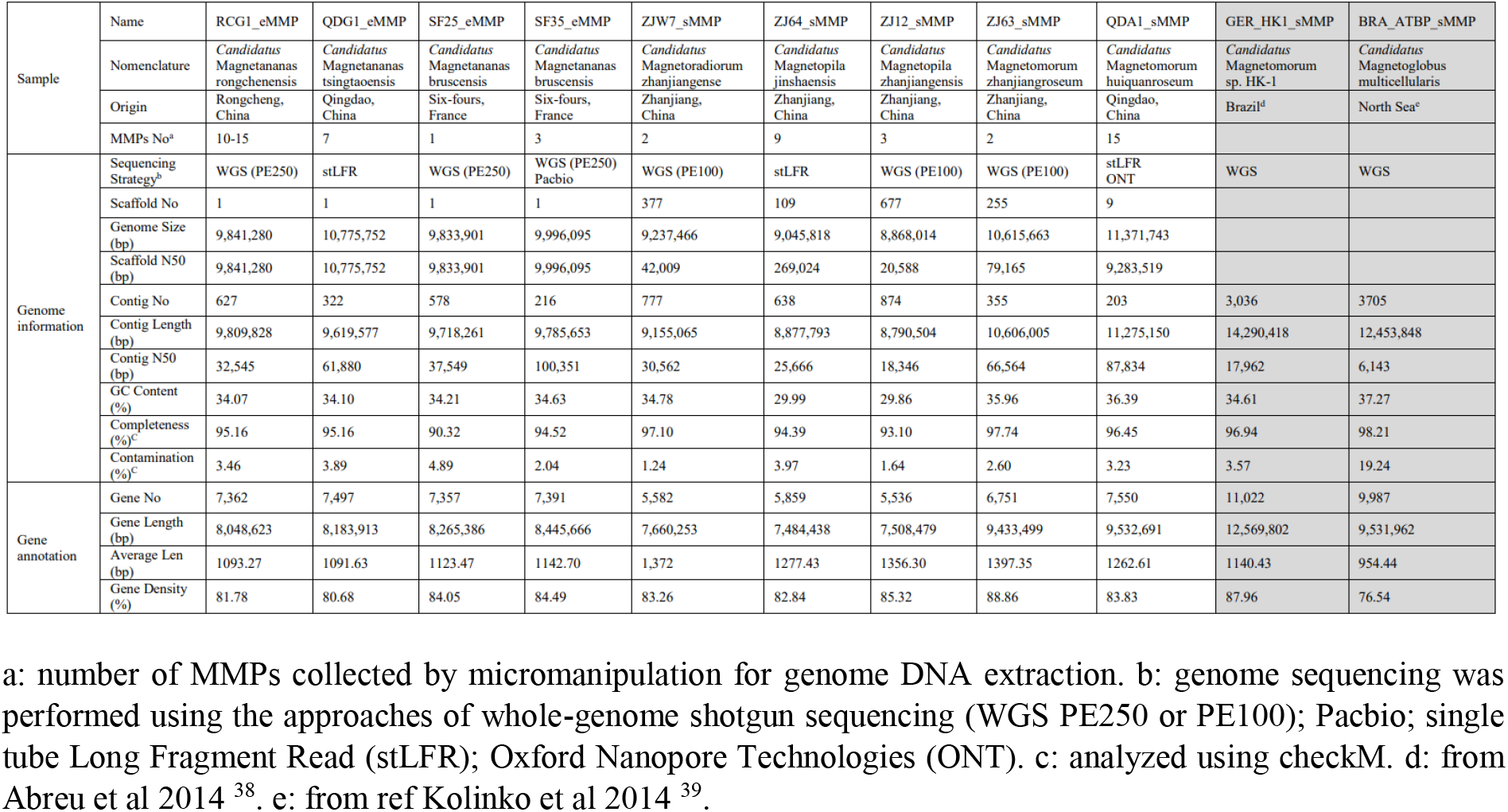
General genomic information of magnetoglobules

We analyzed the evolution of MTBs using 120 bacterial single-copy proteins in the nine magnetoglobule genomes sequenced here, forty-four unicellular MTB and two sMMP genomes available in GTDB. As shown in Figure 3, genomes from Ombitrophus and Nitrospira were clustered together as a branch connected to the Proteobacteria lineage. The large lineage of Proteobacteria was composed of two distinct clades. One contained alphaproteobacterial MTB, etaproteobacterial MTB and gammaproteobacterial MTB subclades. The other had a deltaproteobacterial origin containing the genomes of both unicellular and multicellular MTB. Four unicellular *Desulfovibrio* sp. (IFRC170, MBC34, RS-1 and UBA700) formed the subclade that was removed from the unicellular *Desulfamplus magnetovallimortis* BW-1 and all magnetoglobules. Notably, spherical and ellipsoidal magnetoglobules were clustered into the subclade of magnetoglobule. Among all available genomes in GTDB, the non-magnetic multicellular filamentous bacterium *Desulfonema ischimotonii* Tokyo 01^T^ was the most closely related to magnetoglobules (Figure 3). They clustered together as a multicellular deltaproteobacterial subclade that was linked to a unicellular subclade consisting of unicellular MTB *D. magnetovallimortis* BW-1 and non-magnetic *Desulfobacterium autotrophicum* HRM2. This result indicates that a unicellular deltaproteobacterial ancestor (Figure 3, red asterisk) diverged into unicellular and multicellular deltaproteobacteria. The multicellular branch evolved in non-magnetic filamentous *D. ischimotonii* and magnetotactic magnetoglobules that further diverged to spherical and ellipsoidal magnetoglobules.

**Figure 3.**
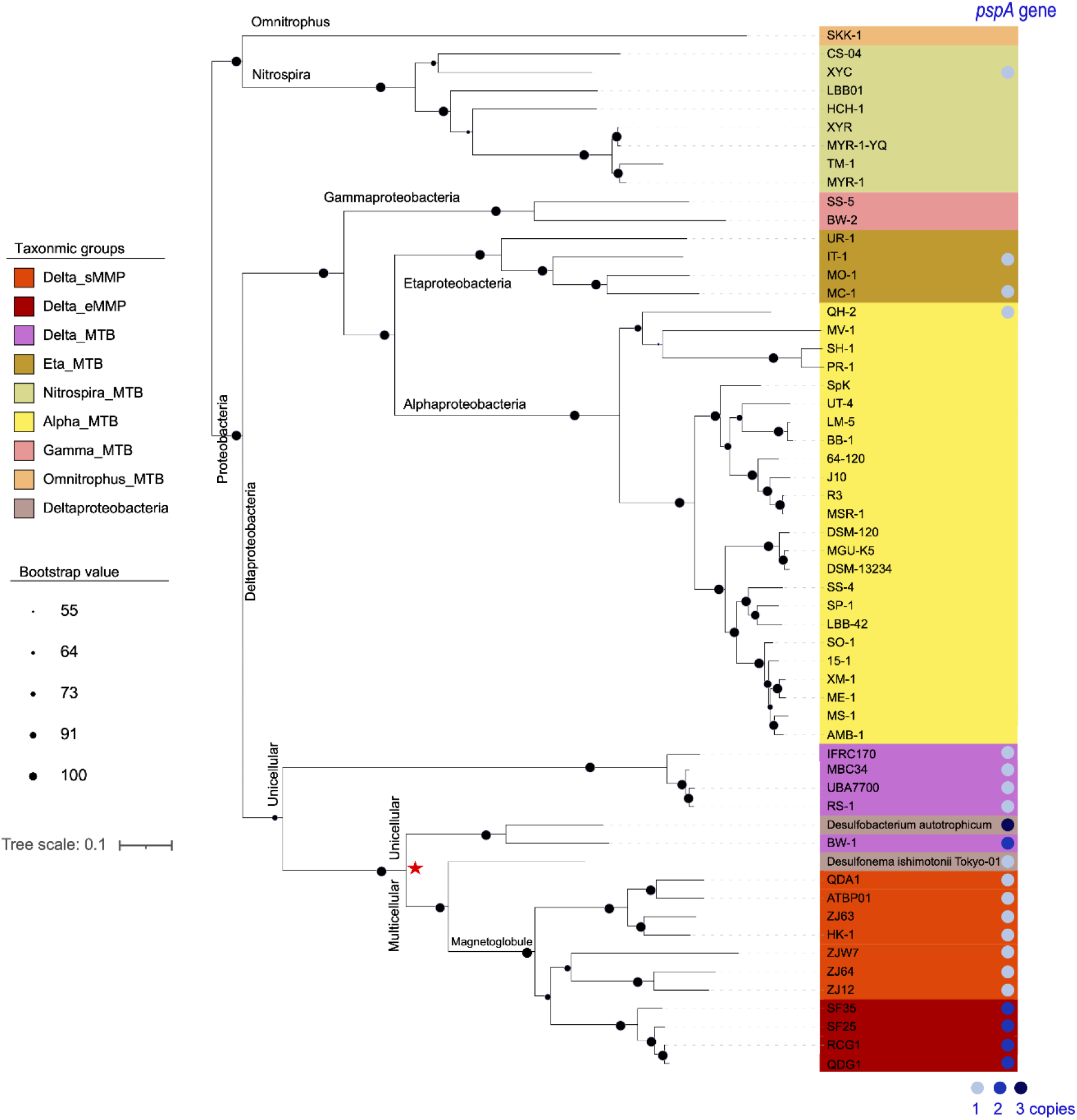
Phylogeny of MTB based on GTDB analysis. Genome phylogenetic tree of 44 unicellular MTB, 11 multicellular magnetoglobules and two non-magnetotactic Desulfobacteraceae genomes based on 120 bacterial single-copy proteins with high branch support values. The red asterisk suggests an ancestor for the unicellular and multicellular magnetotactic bacteria. Distribution of *pspA* in MTB is indicated on right beside the access or species names. Light-blue circle, blue circle and dark-blue circle mean, respectively, 1, 2 or 3 copies of *pspA* detected in the corresponding genome whereas no circle means no *pspA* found in the genome.

To determine the magnetotactic origin of magnetoglobules we studied the phylogeny of *mamB* and *mamM* genes that are essential for the membrane biogenesis of magnetosomes ^14,15,17,18^. Both MamB and MamM clustered together at class level in consistence with their taxonomic affiliation (Figure 4). MamB and MamM phylogenetic trees of deltaproteobacterial MTB both consisted of a magnetite specific clade and a greigite-related clade, which diverged from a common ancestor of deltaproteobacterial MTB. Notably, the two proteins of multicellular magnetoglobules were clustered together with those from the unicellular MTB. Therefore, magnetoglobules acquired magnetosome genes before divergence between magnetite and greigite magnetosomes and shared the same origin with unicellular MTBs. It is likely that magnetoglobules evolved from a unicellular magnetotactic bacteria.

**Figure 4.**
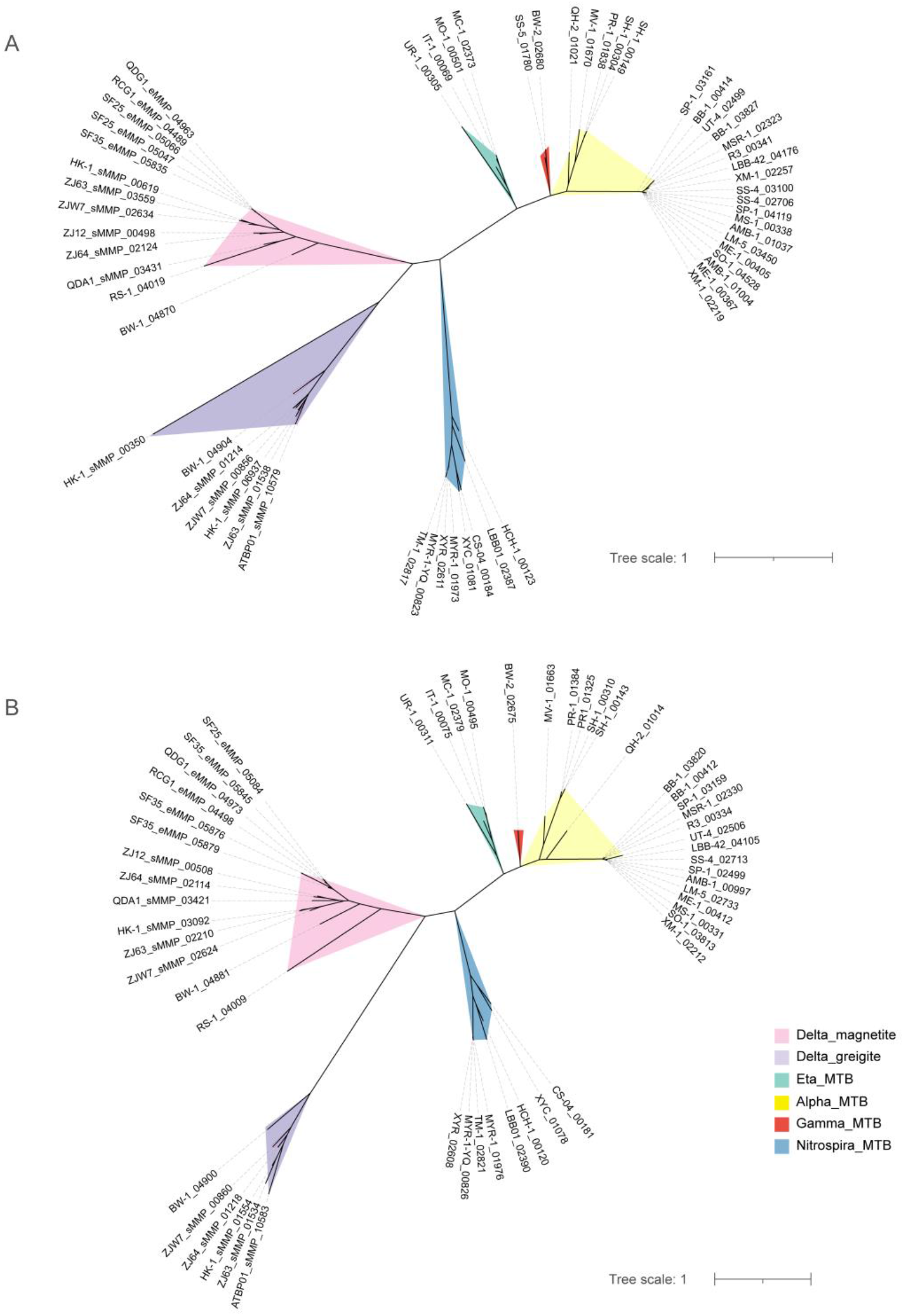
Phylogeny of *mam* genes essential for magnetosome membrane biogenesis. The phylogenetic tree of MamB (A) and MamM (B) proteins from unicellular MTB and multicellular magnetoglobules was constructed using iq-tree software based on their alignment using clustalW2 software, respectively. The taxonomic and magnetosome composition groups were shown with pink (Delta_magnetite), purple (Delta_greigite), green (Eta_MTB), yellow (Alpha_MTB), red (Gamma_MTB), or blue (Nitrospira_MTB) color.

### ESCRT-III proteins are not required for magnetosome biogenesis

Analysis of MTB genomes showed that all deltaproteobacterial MTB had the *pspA* gene (Figure 3). Moreover, the genomes of the unicellular *D. magnetovallimortis* BW-1 and all ellipsoidal magnetoglobules had two copies of *pspA*. Half etaproteobacterial MTB and only one of the 24 alphaproteobacterial MTB or one of the 8 nitrospirial MTB had the *pspA* gene (Figure 3). The two gammaproteobacterial and the omnitrophus MTB had no *pspA*. Non-magnetotactic *D. autotrophicum* HRM2 and *D. ishimotonii* Tokyo 01^T^ had three and one copies of *pspA*, respectively (Figure 3). The large genome size (5.6 Mbp) and high number of genome plasticity elements (> 100 transposon-related genes) might explain the high copy number of *pspA* in strain HRM2. Among the most extensively studied MTB, the alphaproteobacterial MTB *M. magnetotacticum* AMB-1 and *M. gryphiswaldense* MSR-1 had no *pspA*, but the deltaproteobacterial MTB *D. magneticus* RS-1 had it. Therefore, distribution of *pspA* is variable in different taxa, and the membrane-remodeling protein PspA is unlikely required for the biogenesis of magnetosomes.

### Phylogenetic comparison of MTB PspA with those of other bacteria and archaea

Magnetoglobules exhibit conspicuous morphology distinct from other multicellular bacteria and are rich in intracellular vesicles, open rings and tubular rods (Figure 2). All magnetoglobules have PspA and ellipsoidal magnetoglobules have even two copies. The question is whether the membrane-remodeling protein is involved in vesicle biogenesis and the multicellular morphogenesis. We analyzed the phylogenetic relationship of MTB PspA with those of other bacteria and archaea. Remarkably, magnetoglobule PspA and those of the non-magnetic multicellular filamentous bacterium *D. ischimotonii* Tokyo 01^T^ and unicellular deltaproteobacterium *D. autotrophicum* HRM2 clustered in a distinct clade that was separated from other unicellular MTBs and was especially far away from those of other unicellular deltaproteobacterial MTB, except one of the two copies of the unicellular MTB bacterium *D. magnetovallimortis* BW-1 (Figure 5A). More detailed analysis showed that the magnetoglobule clade was composed of PspA of multicellular magnetoglobules, nonmagnetic filamentous *D. ishimotonii* Tokyo 01^T^ and one of the three copies of PspA of *D. autotrophicum* HRM2 (Figure 5B). Notably, the second copy of ellipsoidal magnetoglobule PspA clustered together with relatively weak bootstrap value and far away from other PspA. These results would suggest that *pspA* evolved before occurrence of multicellularity and that it was followed by the *pspA* gene duplication for specialization for ellipsoidal morphogenesis.

**Figure 5.**
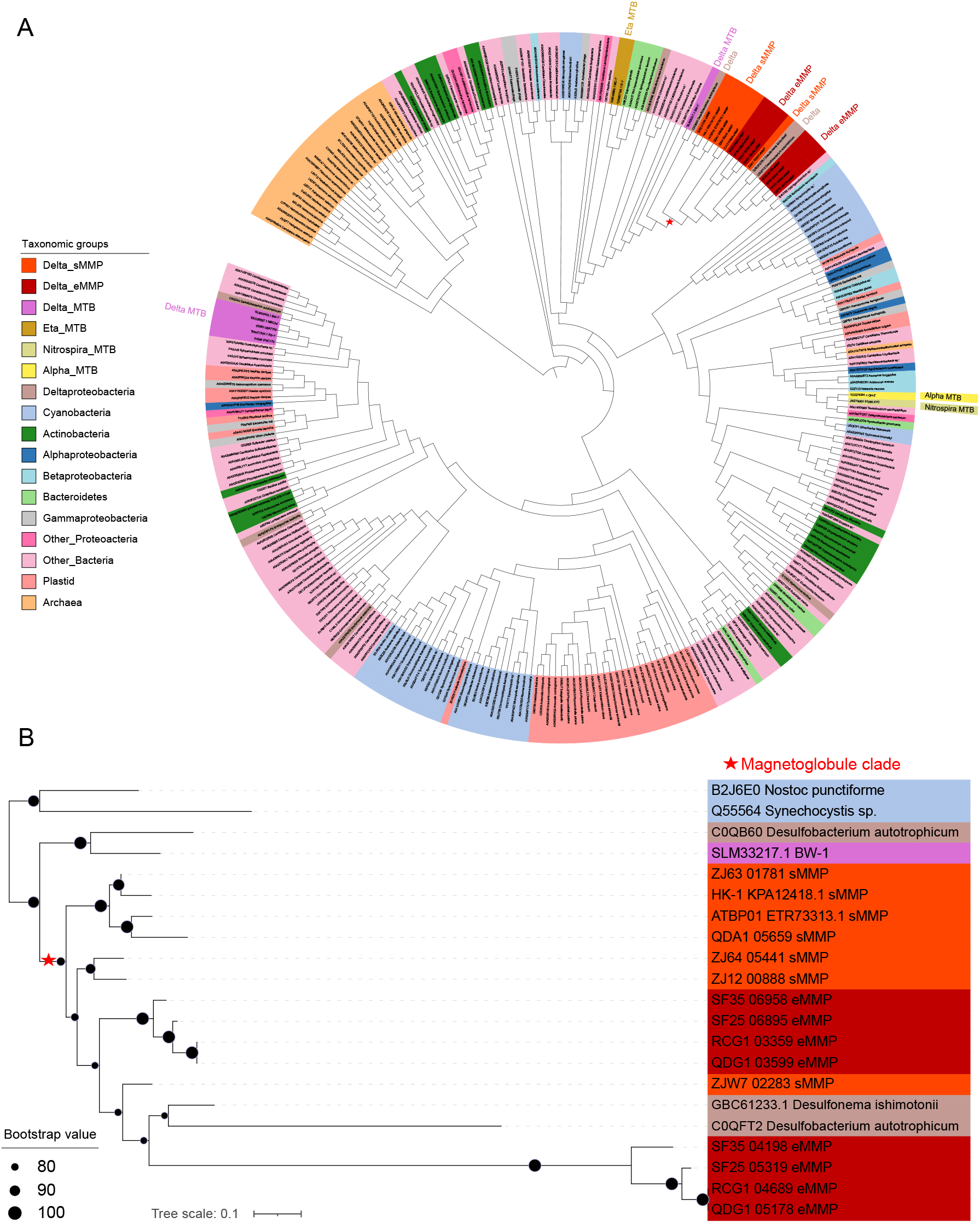
Distribution and phylogenetic analysis of PspA in archaea and bacteria. (A) Phylogenetic tree of 264 representative PspA sequences reported by Liu *et al*. ^3^ (included 26 archaeal, 37 Chloroplast plastid and 201 bacterial sequences) and 27 PspA sequences found in unicellular MTB, multicellular magnetoglobules and the multicellular filamentous *Desulfonema ischimotonii* Tokyo 01^T^. Compared with other bacteria, all the PspA sequences in magnetoglobules formed a distinct clade (red asterisk). (B) More detailed analysis of the magnetoglobule clade. A long branch separates the PspA of *D. ishimotonii* Tokyo 01^T^ and the second copy of eMMPs from the PspA of sMMPs and the first copy of eMMPs. The taxonomic color code is the same as in (A).

### Hetero-expression of ellipsoidal magnetoglobule *pspA* genes in *E. coli* cells

Despite sharing low sequence identity, ESCRT-III proteins exhibit the same secondary structure. Their N-terminal ESCRT-III core domain has four α-helices that fold into a hairpin motif. Additional α-helices (α5 and α6) at the C-terminus might have regulatory function. In the cytoplasm at the free state, the regulatory α5 and α6 fold over the ESCRT-III core domain, in a “closed” conformation, and inhibit polymerization. Interaction of ESCRT-III with the membrane triggers the relief of the auto-inhibition, leading to a conformational change of the proteins into their “open” form and assembly into higher-order structures ^3^. We used AlphaFold to predict the secondary structures and found that the PspA of all spherical magnetoglobules and PspA1 of ellipsoidal magnetoglobules (Figure S1, and as shown by PspA1_eMMP of SF-35 in Figure 6, A1) had similar fold and were composed of additional alpha-helices, thus with a potential regulatory domain. All these PspA monomer exhibited a predicted open conformation as the human ESCRT-III CHMP1B and activated membrane-binding Snf7 ^40^. In contrast, the PspA2 of the four ellipsoidal magnetoglobules (Figure S1, and represented by PspA2_eMMP of SF-35 in Figure 6, A2) had the well conserved α1-α2/α3 hairpin, but exhibited subsequent folding patterns distinct from PspA1. Their Hinge 2 (elbow) bended α4 to form an antiparallel alpha-helix structure with α2/α3, which is similar to the closed conformation of yeast ESCRT-III Snf7 and Vps24 ^40^. Such a structure might require interaction with the membrane or other components to release the inhibition and assemble into a diverse set of flexible polymers that contributes to the architecture and morphology of ellipsoidal magnetoglobules.

**Figure 6.**
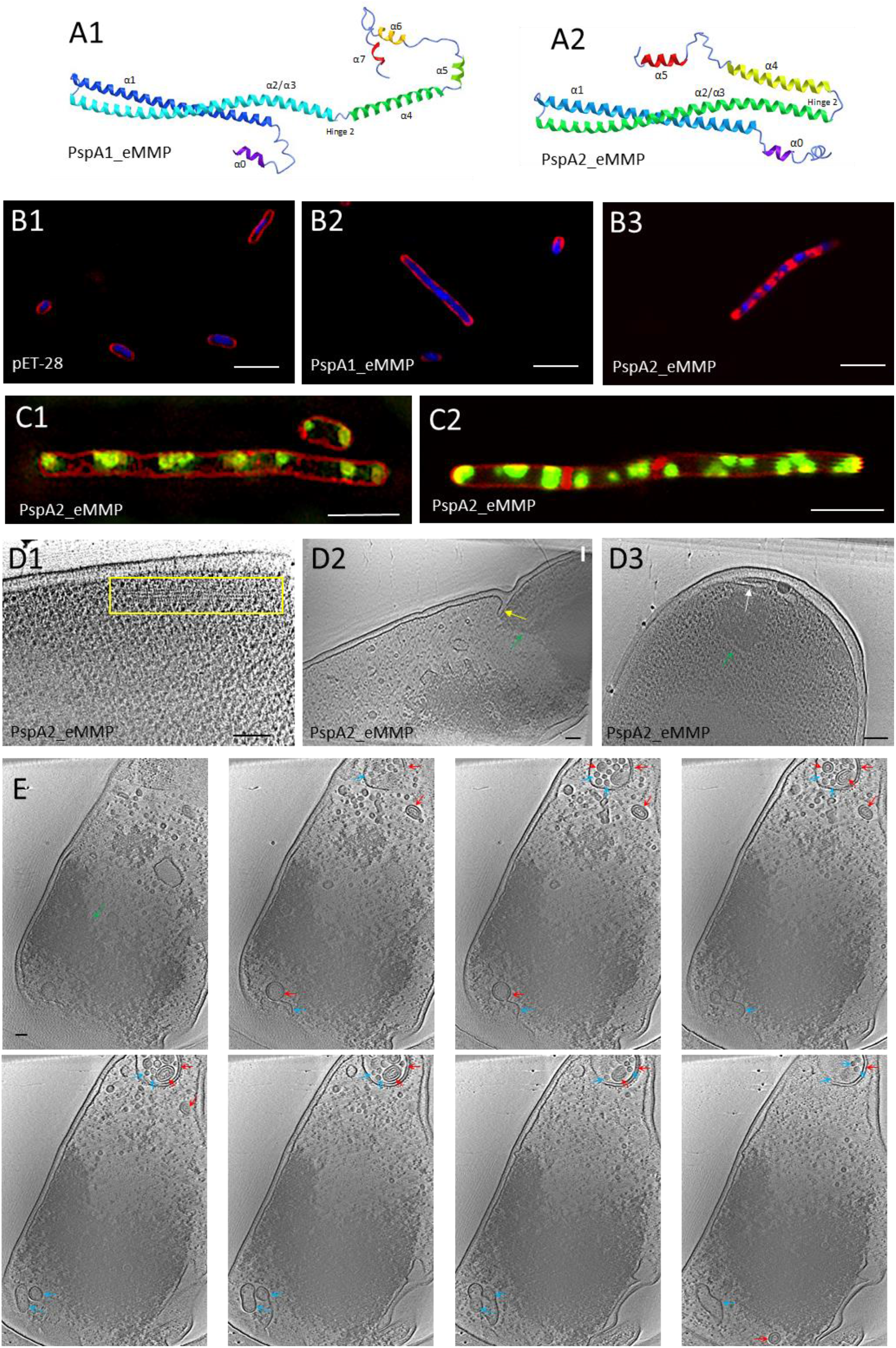
Morphology of *E. coli* cells expressing *pspA* genes of *Candidatus* M. bruscensis SF-35. AlphaFold predicated secondary structure of PspA1 (A1, PspA1_eMMP) and PspA2 (A2, PspA2_eMMP) monomer of SF35. (B) Laser confocal images of *E. coli* cells carrying vector pET-28 (B1) or its derivative plasmids expressing *pspA1* (B2, PspA1_eMMP) or *pspA2* (B3, PspA2_eMMP) of SF35. (C) 3D-SIM (C1) and Laser confocal (C2) microscope images show the polar location of mNeonGreen-PspA2 in unicell and regular distribution in filamentous cells expressing PspA2 fusions. The membrane is stained by FM4-64 (red) and DNA by DAPI (blue). mNeonGreen-PspA2 are in green color. (D) CET micrograph gallery shows filament bundles (D1, yellow square), affected septum (D2, yellow arrow) and membrane sacculus (D3, white arrow). Green arrows indicate electron dense area. Panel E shows tomograph of an *E. coli* cell expressing *pspA2* of SF35 with multivesicular body-like structure and formation of double membrane (red arrows) and single membrane (cyan arrow) vesicles. Scale bars are 5 µm in B and C and 100 nm in D and E.

To assess membrane remodeling capacity of ellipsoidal magnetoglobule PspA proteins, we expressed the *pspA1* and *pspA2* genes of the *Candidatus* M. bruscensis SF-35 in *E. coli*. The 6His-tag and Tev cleavage site were added at the N-terminus of these PspA proteins. IPTG induced the overexpression of the recombined *pspA2* genes with sizes similar to the expected products (∼30 kDa), but increased sizes for PspA1_eMMP (∼ 36 kDa versus 30.8 kDa) (Figure S2, A). Optical microscope analysis showed an increase of the median length of the cells from 3.48 µm for the strain carrying the plasmid vector to 4.21 µm and 4.87 µm for those expressing PspA1_eMMP and PspA2_eMMP, respectively (Figure S2, B). Consistently, laser confocal microscopy (Figure 6, B) and three-dimensional structured-illumination microscopy (3D-SIM) analyses (Figure S3) showed that the vector had no noticeable effect on cell division, but expression of multicellular PspA1_eMMP and PspA2_eMMP resulted in increased long undivided cells (Figure 6, B). Hydrophilic styryl dyes FM4-64 is capable of incorporating in the outer leaflet of the plasma lipid bilayer without penetration through the plasma membrane, and its intercalation into hydrophobic membrane enhances the fluorescence. Therefore, FM4-64 is internalized exclusively by endocytic process and commonly used for the tracking of endocytosis and exocytosis in eukaryotic cells ^41,42^. It is also suitable for fluorescence imaging of magnetoglobule membrane ^30^. In *E. coli* strains carrying the plasmid vector pET-28 or expressing PspA1_eMMP, FM4-64 stained the periphery of cells (Figure 6, B2 and Figure S3). In contrast, FM4-64 clearly aggregated at positions between DAPI stained DNA spots in the cytoplasm of the cells expressing PspA2_eMMP (Figure 6, B3 and Figure S3). The intracellular location of bright red fluorescence indicated intercalated FM4-64 in hydrophobic membrane and implied membrane remodeling in the cells. We then labeled PspA2 by in-frame fusion of mNeonGreen between the Tev and PspA2. 3D-SIM (Figure 6, C1) and laser confocal (Figure 6, C2) imaging showed that PspA2 was located at both poles of rod-shaped unicells and distributed as foci in filamentous cells. Membrane invagination was observed in 3D-SIM micrographs.

### Cryo-electron tomography of *E. coli* cells expressing the *pspA2* gene

Cryo-electron tomography analysis was carried out to investigate the membrane structure of the *E. coli* cells expressing the *pspA2* gene. Filament bundles were observed in some cells (Figure 6, D1). In long filamentous cells, electron dense area was separated by electron weak area (Figure S4, Figure 6, D and E, green arrows), which was reminiscent to the patterns of regular distribution of mNeonGreen-PspA2 spots as observed in laser confocal and 3D-SIM micrographs (Figure 6, C). Further detailed inspection of the tilt series and reconstruct them into 3D tomogram revealed that some electron dense, aggregate-like area at septum impaired the inward orientation of the constriction ring (Figure 6, D2). Invagination of inner membrane into cytoplasm as saccules was evident (Figure 6, D3, Movie S1). In addition, unilamellar and bilamellar vesicles were generated, and some of them were enclosed in a double membrane compartment that was reminiscent to multivesicular body or compartment of unconventional protein secretion (Figure 6, E). These structures imply that expression of the *pspA2* gene of *Candidatus* M. bruscensis SF-35 alone is sufficient to result in membrane remodeling in *E. coli* cells.

## Discussion

Expansions of genome size during evolution seems to be a strategy used by bacteria for their adaptation to various environments. Myxobacteria are another multicellular deltaproteobacterial taxon dwelling in soils, river mud, deep-sea sediments and hydrothermal vents ^43^. Sequencing of the largest genome (16.04 Mbp) of myxobacterium *Minisystis rosea* DSM 24000^T^ and comparative genomics analysis of 22 myxobacterial genomes, ranging from 4.35 Mbp to 16.04 Mbp with a median size of 11.53 Mbp, showed a strong positive correlation between genome size and number of genes involved in signal transduction or coding for secretome proteins ^44^. Multicellular magnetoglobules had approximate double sizes (average: 10.6 Mbp, n=11) compared to unicellular deltaproteobacterial MTB (average: 5.1 Mbp, n=5). Compared to the unicellular deltaproteobacterial MTBs, magnetoglobules COGs (Cluster of Orthologous Groups of proteins) were significantly expanded in T (Signal transduction mechanisms), U (Intracellular trafficking, secretion, and vesicular transport), O (Posttranslational modification, protein turnover, chaperones), Q (Secondary metabolites biosynthesis, transport and catabolism), K (Transcription), N (Cell motility), L (Replication, recombination and repair) and D (Cell wall/membrane/envelope biogenesis), as well as that of general function prediction only class (R) and function unknown class (S). On the contrary, contraction was obvious in some classes related to basic metabolism, such as P (Inorganic ion transport and metabolism), C (Energy production and conversion) and E (Amino acid transport and metabolism). Notably, multicellular magnetoglobules encoded more genes involved in signal transduction and transcription, which are particularly relevant for multicellularity as reported ^45,46^. Gene family clustering analysis also revealed a significant increase (greater than 5) of 58 orthologous proteins in magnetoglobules compared to unicellular deltaproteobacterial MTB. Among them 9 orthologous proteins were classified as filamentous haemagglutinin, cadherin, cadherin-like, cellulosome anchoring protein, Na-Ca exchanger/integrin-beta4, and fibronectin type III. These proteins might be responsible for cell adhesion and contributed to morphogenesis and development of multicellular organization.

Multicellular magnetoglobules might evolved from a non-magnetic unicellular ancestor to multicellular organisms and then obtained the magnetosome gene cluster (MGC) via horizontal gene transfer. Lefèvre *et al*. have collected non-magnetotactic MMPs (nMMP) from low-saline, nonmarine environments ^47^. This report supports the hypothesis of emergency of magnetoglobules from unicellular non-magnetic bacterium via an intermediate stage of nMMP. Alternatively, the ancestor of magnetoglobules was a unicellular MTB that evolved to multicellular magnetoglobules. During the evolution some of the descendants lost MGC and became non-magnetotactic, probably as in the case of nMMP, as well as the unicellular *D. autotrophicum* HRM2 and filamentous *D. ischimotonii* Tokyo 01^T^ that are the most closely related to magnetoglobules. The fact that the phylogenetic positions of the *mamB* and *mamM* genes of magnetoglobules correspond to those on the genomic taxonomic tree indicates a vertical inheritance and evolution of MGC in magnetoglobules, which is in favor for the second hypothesis. Based on the phylogenetic results of whole genome and PspA, we propose the evolution of magnetoglobules from a unicellular magnetotactic deltaproteobacterium ancestor.

Intracellular compartmentation is a key step in the evolution of multicellularity and eukaryotic organisms. Development of intra- and intercellular compartments and exchange of nutrients and information among these compartments require extensive membrane remodeling. It has been reported that 19 types of bacterial compartments were observed in at least 23 phyla ^48-50^. These compartments have been defined as a proteomically defined lumen bound by a lipid bilayer (membrane), a lipid monolayer, a proteinaceous coat or phase-defined boundary ^48^. Besides the magnetosomes, magnetoglobules contain vesicles, lipid inclusions and polyphosphate granules, which can be also considered as cellular compartments. The membrane remodeling proteins might be required for generation of the membrane enclosed compartments such as thylakoids and magnetosomes. The requirement for the membrane remodeling proteins PspA, Imm30 and Vippp1 for thylakoid biogenesis has been well documented ^3,5,6,34,51^. Our phylogeny analysis and experimental results reported by others rule out the requirement of PspA for the formation of magnetosomes ^12-15,17,18^. Interestingly all magnetoglobules have PspA that form a detached clade and ellipsoidal magnetoglobules have the duplicated PspA2 paralogs. Consistent with the presence of these membrane-remodeling proteins, vesicles were found in the core lumen, in the out-surface matrix, and appeared as multivesicular body in the periplasm of cells (Figure 2). In addition, filaments, tubular rods and open ring structures were abundant in the cytoplasm of these cells. Intriguingly how the vesicles are severed in magnetoglobules that don’t have counterpart of the Vps4 ATPase. Nevertheless, hetero-expression of magnetoglobule *pspA2* genes in *E. coli* resulted in the formation of vesicles, multivesicular body and filaments similar to those observed in magnetoglobule cells (Figure 6 versus Figure 2), showing a membrane remodeling capacity.

Division mechanism of Archaea is more complex than that of bacteria. Euryarchaeota possess FtsZ homologs and divide via an FtsZ-based mechanism that is similar to the bacterial division ^52^. Interestingly, in some orders of Crenarchaeota and the Asgard super-phylum of Archaea, cell division depends on Cdv (for cell division) machinery that consists of *cdvABC* ^53,54^. CdvA is found only in archaea whereas CdvB and CdvC are homologous to the eukaryotic ESCRT-III and Vps4 (vacuolar protein sorting), respectively ^54^. During Cdv-based division process, CdvA is targeted to the division site and recruit the other two Cdv components. The CdvB forms a contractile machinery to sever the membrane neck. CdvB are classified into Vps2/24/46 class and Vps20/32/60 class ^54^. The CdvC, eukaryotic Vps4 homologous ATPase, interacts with MIM1 motif (leucine-rich motifs in Asgard) in the C-terminal helix of the Vps2/24/46 class ESCRT-III subunits, and MIM2 motif (proline-rich motifs in Asgard) in the C-terminus of Vps20/32/60 class subunits to disassemble the membrane abscission polymers and turnover CdvB. The Cdv proteins function in exovesicle secretion, viral release, and cell division ^54^. *Sulfolobus islandicus* REY15A has three ESRCT-III paralogs. Liu et al. have shown that ESCRT-III, ESCRT-III-1 and ESCRT-III-2 play a crucial role at different stages of membrane ingression during cell division and ESCRT-III-3 is essential for cell budding ^55^. In Sulfolobales without CdvA, CdvB and CdvC are sufficient to generate exovesicles ^54^. Magnetoglobule genomes encode only CdvB homologs, PspA1 and PspA2, with neither CdvA nor CdvC counterparts. Consistently, their PspAs didn’t have MIM1 or MIM2 motifs (Figure S5). All magnetoglobules possess the complete FtsZ and Min machineries. Therefore, magnetoglobule cells can divide via the canonical bacterial FtsZ-based mechanism. Whether the PspAs are involved in magnetoglobule cell division is unknown. The ESCRT of budding yeast serves to mediate the turnover of cell-division proteins from the plasma membrane ^56^, or to the control of membrane trafficking during cytokinesis ^57^ instead of a direct action in abscission. The PspA might be required essentially for the vesicle formation and probably is involved in the morphogenesis of ellipsoidal magnetoglobules. Synthetic biology analysis could partially circumvent the problem of missing of magnetoglobule cultures in determining the function of the membrane remodeling PspA proteins. High resolution structures of PspA homo- and hetero-polymers, both in vivo/situ in magnetoglobules and in synthetic cells or in vitro in various solutions with or without lipid, could shed light on the function of magnetoglobule PspA.

Bacterial PspA was first discovered in *E. coli* ^58^. It is encoded by *pspABCDE* operon that is transcribed in opposite direction compared to that of the upstream *pspF* gene. *E. coli* PspA has dual functions. It forms PspA-PspF complex and blocks the activation of *pspABCDE* transcription by the enhancer-binding protein PspF. When phage pore-forming protein secretin pIV assembles in outer-membrane, the resulting membrane stress triggers the interaction of PspA with the membrane PspB-PspC complex and inner membrane, which releases PspF and activates the transcription of *pspABCDE*. PspA oligomers stabilize compromised areas of the inner membrane and maintains the membrane integrity. Other factors perturbing membrane such as mislocalization of secretin of protein secretion systems, impairment of protein export and environmental extremes can also enhance PspA production ^59^. Magnetoglobules have only PspA, without PspBCDE components. However, they do possess type I, II, IV and VI secretion systems and corresponding secretin, e.g. PilQ. Therefore, PspA might play a role in maintaining the membrane integrity in magnetoglobules without PspB and PspC. Interestingly, pIV secretin-dependent induction of the Psp response and activation of the Psp response by heat shock has also been reported as independent of PspB and PspC in *E. coli* ^59,60^.

In *E. coli*, PspA seems to be functionally linked with the cytoskeleton proteins MreB and RodZ that maintain rod-shape cellular morphology ^61-63^. These results imply existence of unrecognized relationships between PspA membrane remodeling function and multicellular morphogenesis. Our phylogenetic analysis showed that most PspA from multicellular cyanobacteria trend to cluster in clades separated from those of mainly unicellular cyanobacteria (Figure S6). We initiated studies to evaluate morphogenesis function of PspA by hetero-expression of magnetoglobule PspA1 and PspA2 in *E. coli*. Their expression could result in filamentous morphotype of *E. coli*. In addition, hetero-expression of PspA2 led to the formation of multivesicular body-like structures, vesicles and filament bundles. Further detailed studies are needed to identify the protein composition of these compartments and elucidate the mechanism of interference with cell division and separation. According to their in vivo function, ESCRT-III can be classified as essential, helpers or special membrane remodeling proteins ^54^. PspA1 are clustered with other PspA whereas PspA2 form a separated clade. Therefore, duplication of PspA1 to create the second copy of PspA2 might be the requirement of the ellipsoidal morphogenesis.

## Methods

### Sample collection and whole genome application

Samples were collected from Yuehu Lake, Rongcheng city (RCG1), Huiquan Bay, Qingdao city (QDG1 and QDA1), Brusc lagoon, Six-Four les Plages, Southern France (SF25 and SF35) and Jinsha Bay, Zhanjiang city (ZJ64, ZJ12, ZJW7 and ZJ63). Magnetoglobules were micro-sorted using a TransferMan ONM-2D micromanipulator and a CellTram Oil manual hydraulic pressure-control system (IM-9B) equipped on a microscope (Olympus IX51), after magnetic enrichment from the Intertidal sediment ^39,64-66^. One to fifteen micro-sorted magnetoglobules were stored in PBS, then whole genome amplification (WGA) was performed using the multiple displacement amplification (MDA) with REPLI-g Single Cell kit, according to the manufacturer’s instructions ^25^.

### Genome sequencing, assembly and annotation

The WGA products were prepared for library construction using strategy of whole-genome shotgun sequencing (WGS), single tube Long Fragment Read sequencing (stLFR), respectively. The WGS libraries of RCG1, SF25 and SF35 were sequenced on Illumina MiSeq platform (BGI-Wuhan, China) to generate 250 bp paired-end raw reads. The WGS libraries of ZJ12, ZJW7, ZJ63 and stLFR libraries of QDG1, ZJ64, QDA1 were sequenced on BGISEQ-500 platform (BGI-Qingdao, China) in 100 bp pair-end model. After performing quality trimming and filtering using SOAPnuke, the high-quality clean reads of WGS data were assembled using metaSPAdes (v3.14.1) with k-mer size from 33 to 113 by step 20, and the stLFR clean reads were assembled using Supernova (v2.1.1) as described before ^67^.

For all primary assemblies, the MetaWRAP pipeline was performed for the metagenome binning by using “-metabat2 -maxbin -concoct” modules, and then using “bin_refinement” and “reassemble_bins” modules to generate the high-quality recovered genomes ^68^. The genome quality was evaluated by using CheckM ^69^, and the magnetoglobule draft genomes with lower contamination (<5%) and higher completeness (>90%) were obtained.

To construct the longer contiguity genomes, SLR-superscaffolder (v0.9.0) ^70^ was applied for the QDG1, ZJ64 and QDA1 genome scaffolding. Meanwhile, Oxford Nanopore Technologies (ONT) and PacBio continuous long read (CLR) sequencing were used for QDA1 and SF25 respectively, and then SSPACE-LongRead (v1.1) and TGS-Gapcloser (v1.1.1) ^71^ were applied for genome scaffolding and gap closed. Furthermore, RaGOO (v1.1) was used for the genome scaffolding of the magnetoglobule genomes within the same genus, and GMcloser (v1.6) was used for gap closed by using WGS or stLFR sequencing reads with parameters “-mm 500 -mi 95 -ms 1000 -l 300 -i 400 -d 100”, then Pilon (v 1.23) and GATK (v 3.4-0) were used to fix the sequencing errors. Finally, four ellipsoidal magnetoglobule pseudo-chromosomal level genomes and five near complete spherical magnetoglobule genomes were obtained.

The general information of genomes were stated using Quast ^72^, genes were predicted and annotated using Prokka ^73^ and Microbial Genome Annotation and Analysis Platform (MaGe, https://mage.genoscope.cns.fr/microscope) ^74^. 16S rRNA identity and average nucleotide identity (ANI) were calculated by blast and FastANI, respectively. Sequencing, assembly and quality control flow was present in Supplementary information (Figure S7).

### Genome features and comparative genomic analyses

A concatenated alignment of the 120 bacterial single-copy proteins of the 9 magnetoglobules genomes together with the two public spherical magnetoglobules genomes, 44 unicellular MTB and 2 Deltaproteobacterial non-magnetic bacteria were analyzed using GTDB-tk (v.1.7.0) based on Genome Taxonomy Database (GTDB, release 202) ^75,76^, then maximum likelihood (ML) genome phylogenetic tree was constructed using IQ-Tree (v2.0.4) ^77^, and bootstrap values were calculated with 1000 replicates under the LG+R6 model test by -IQ-Tree with Bayesian information criterion (BIC). Magnetosome genes of magnetoglobules genomes were identified using the MagCluster ^78^. The phylogenetic trees based on magnetosome proteins MamB and MamM were constructed by ML method using the software IQ-Tree. The *pspA* genes of magnetoglobules and unicellular MTB genomes, were searched in Pfam database with E-valule < 1e-5 by using InterProScan (v5), and the phylogeny was inferred by using IQ-Tree under the best-fitting LG+C30+G+F model. All the phylogenetic trees were visualized and adjusted using iTOL (https://itol.embl.de/). The secondary structures of PspA sequences were predicted by AlphaFold2 using the pdb70 template mode ^79,80^.

### Molecular and protein analyses

The *pspA1* and *pspA2* genes of SF35 were synthesized and cloned in pET-28a(+)-Tev plasmid (GenScript Biotech (Netherlands) BV), which were transformed in *E. coli* strain BL21(DE3). Using InFusion procedure, mNeonGreen gene was in-frame inserted between the Tev and PspA2 and transformed in BL21(DE3). Transformants were grown in LB media to OD_600nm_ = 0.4 to 0.6, and expression of *pspA* was induced by adding isopropylthio-β-D-galactoside (IPTG) to a final concentration of 0.5 or 2 μM. Four hours after the induction the cells were harvest by centrifugation, washed with PBS buffer, and used for microscopy or biochemistry analyses. Cells were broken by lyse-loading buffer and analyzed on 10% SDS-PAGE.

### Microscopy analyses

Routine optical microscopy observation was performed with Zeiss Axiostar Plus, Zeiss Axio Vert 200M, and Olympus BX51. To perform laser confocal analysis, cells were fixed with 4% paraformaldehyde for 2 hours at room temperature or overnight at 4℃, stained with 7.5 µg/ml FM4-64 (for membranes) and 2 µg/ml DAPI (for chromosomal DNA) and observed with the Olympus FV1000 microscope with laser 405 nm excitation and 425–475 nm emission for DAPI and 543 nm excitation and 555–655 nm emission for FM4-64. Images were collected at a series of focal levels at 0.17 µm intervals.

3D-SIM was performed on a microscope system (DeltaVision OMX SR, GE Healthcare UK Ltd). Images were acquired using a Plan Apo N × 60, 1.42 NA oil immersion objective lens (Olympus, Tokyo, Japan) and two liquid-cooled sCMOs cameras (PCO, Kelheim, Germany). Imaging was performed using excitation at 488 nm during 10 ms at 50%T for mNeonGreen, and excitation at 568 nm during 50 ms at 30%T for FM4-64 dye. Fluorescence was respectively recovered with a 528/48 nm emission filter and a 609/37 nm emission filter. 3D-SIM images were realized by acquiring a z-stack of 11 images separated by 0.125 μm. For each z-section 15 images (5 phases and 3 rotations) were acquired. Images were then reconstructed using the DeltaVision OMX SoftWoRx 7.0 software package (GE Healthcare). The resulting size of the reconstructed images was of 1024 × 1024 pixels from an initial set of 512 × 512 raw images. The channels were then carefully aligned using alignment parameters from control measurements with Image registration calibration slide and 0.1 μm TetraSpeck™ Fluorescent Microspheres (Molecular Probes, Eugene, OR, USA).

### Electron Microscopy

Magnetotactic bacteria were pelleted, high-pressure frozen, freeze substituted, embedded in Epon resin (Medium Grade). Preparation of HPF/FS ultra-thin sections (60–90 nm) was the same as previously described in ^30^. The samples were analyzed using a Tecnai 200 kV electron microscope (FEI) and digital acquisitions were made using a numeric camera (Eagle, FEI).

### HAADF-STEM elemental composition analysis

Magnetoglobule granules were investigated using a JEM-2100F microscope (JEOL Ltd) operating at 200 kV equipped with a Schottky emitter, an ultra-high resolution (UHR) pole piece and an X-ray energy dispersive spectrometer (XEDS). HAADF-STEM was used for Z-contrast imaging. Chemical compositional analysis was performed by STEM-XEDS elemental mapping.

### Cryo-electron tomography

A 5 µl cell suspension containing 10 nm colloidal gold particles were deposited on the copper EM grids with holey carbon supported film (Quantifoil, 200mesh, R1/2). The EM grid were blotted with filter paper (Waterman, grade 1) and plunge-frozen in liquid ethane using gravity-driven plunger apparatus. The grids were transferred into the dewar filled with liquid nitrogen for the storage.

The tilt series were collected using a Titan Krios microscope equipped with 300 kV field emission gun and K3 summit direct detection camera. The images were collected at focus using a volta phase plate and energy filter with a 20-eV slit. SerialEM software were used to collect tilt series images ^81^. The magnification of 26,000 was used and resulted in a physical resolution of 0.338 nm/pixel. The cumulative dose is 100e/ Å, distributed 35 tilt series images taken by dose symmetric scheme. The tilt angles are ranging from −51° to +51° at a step size of 3°. The initial starting tilt is collected at 0°. For every single tilt series collection, the dose-fractionated movie mode was used to generate 8 sub-frames per projection image. Collected dose-fractionated data were first subjected to the motion correction program to generate drift-corrected stack files ^82^. The stack files were aligned using gold fiducial markers and volumes reconstructed by the weighted back-projection method, using IMOD ^83^.

## Data availability

The 9 magnetoglobule assembled genomes and the PspA, MamB and mamM gene sequences analyzed in this study have been deposited into China National GeneBank Sequence Archive (CNSA, https://db.cngb.org/cnsa/) of China National GeneBank DataBase (CNGBdb) with accession number CNP0003599.

## Acknowledgements

We thank A. Bernadac for critical reading of the manuscript, F. Alberto for suggestion for *pspA* cloning, L. Espinosa pour fluorescence microscope assistance, H.M. Yang and T. Jin for Initiation and continuous support of this study. This work was supported by a funding from the Excellence Initiative of Aix-Marseille University -A*Midex, a French “Investissements d’Avenir” programme, by a grant DBM2021 from CNRS, by grants U1706208, 41330962 from NSFC and grants from CNRS for LIA-MagMC. The LABGeM (CEA/Genoscope & CNRS UMR8030), the France Génomique and French Bioinformatics Institute national infrastructures (funded as part of Investissement d’Avenir program managed by Agence Nationale pour la Recherche, contracts ANR-10-INBS-09 and ANR-11-INBS-0013) are acknowledged for support within the MicroScope annotation platform.

## Author contributions

LF.W. designed and LF.W., T.X., W.Z. and J.C. led the project. C.S., XX.Q, LF.W., HM.P., YR.C., J.L., YC.Z. and KX.C., collected the samples. W.Z., YR.C., J.L., YC.Z. and KX.C. prepared DNA for sequencing. W.Z., J.C. and GL.L. conducted genomic analysis. A.K., H.LG., J.D., X.L., N.M. and E.D. performed microscopy analyses. S.Z. carried out CET analysis. J.D. and X.L. constructed the *pspA* expression strains. LF.W., J.C. and W.Z wrote the manuscript. C.S., T.X., HM.P., WJ.Z., A.R. revised the paper. W.Z, J.C. and J.D. contributed equally.

