## supplementary information for "Membrane-remodeling protein ESCRT-III homologs incarnate the evolution and morphogenesis of multicellular magnetotactic bacteria"

### contribute equally.

This file contains:

Figure S1 AlphaFold predicated secondary structure of PspA monomer of magnetoglobules.

Figure S2. Expression of representative *pspA* genes in *E. coli.*

Figure S3. 3D-SIM image of *E coli* expressing PspA1 and PspA2 of ellipsoidal magnetoglobule SF35.

Figure S4. CET analysis of intracellular structures of *E. coli* strain expressing *pspA.*

Figure S5. Multiple alignment of PspA proteins.

Figure S6. Phylogeny of cyanobacteria PspA.

Figure S7. Sequence, assembly and genome recovery flow chart of the study.

Table S1. Pairwise comparison of 16S rRNA genes and average nucleotide identity (ANI).

Movie S1. PspA2 mediated formation of membrane sacculus.

Cryo-electron tomography of a *E. coli* cell expressing ellipsoidal magnetoglobule *pspA2* gene.

Data S1. Metadata summary of analyzed genomes, pspA, MamB and mamM in Figures 3, 4 and 5.

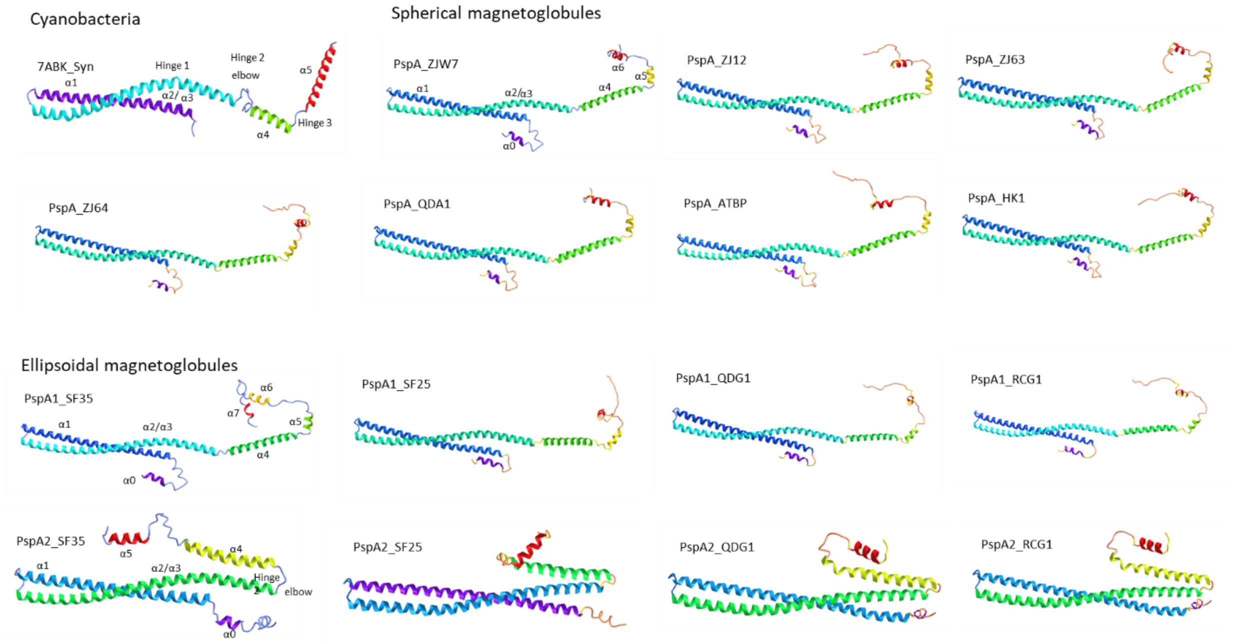


Figure S1. AlphaFold predicated secondary structure of PspA monomer of magnetoglobules compared with cyanobacterium PspA. The structures were drawn using icn3d at https://www.ncbi.nlm.nih.gov/Structure/icn3d/full.html. The secondary structures are shown with spectrum color from N- to C-termini (purple to red). 7ABK: Synechocystis PspA. Magnetoblobule PspA were named according to their genome names.

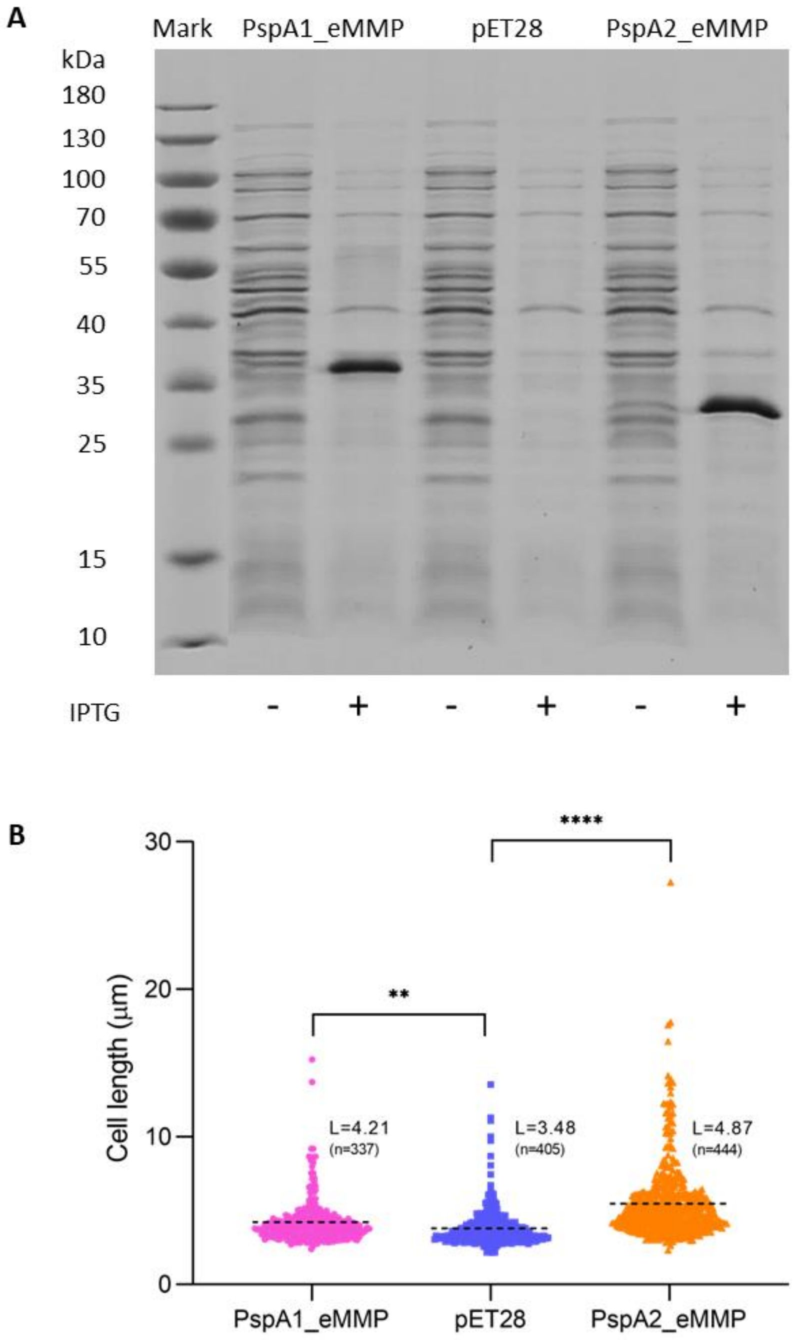


Figure S2. Hetero-expression of the *pspA* genes in *E. coli* cells. (A) Crude extracts of *E. coli* BL21 cells carrying pET28, PspA1_eMMP and PspA2_eMMP were resolved on SDS-PAGE and polypeptides were visualized by Coomassie blue staining. (B) Median cell length measurement of the 3 cultures. n=405, 337 and 444 biologically independent cells in the three cultures, respectively. L represents the median cell length (dashed line). Statistical test used unpaired t test (*****p* < 0.0001, ***p* < 0.005).

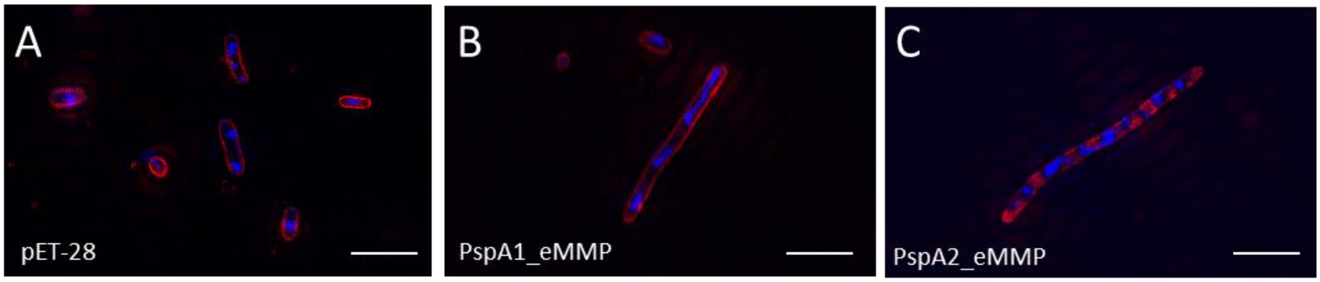


Figure S3. 3D-SIM image of *E coli* expressing PspA1 (B) and PspA2 (C) of ellipsoidal magnetoglobule SF35 compared to the cells with vector plasmid in (A), scale bars are 5 μm.

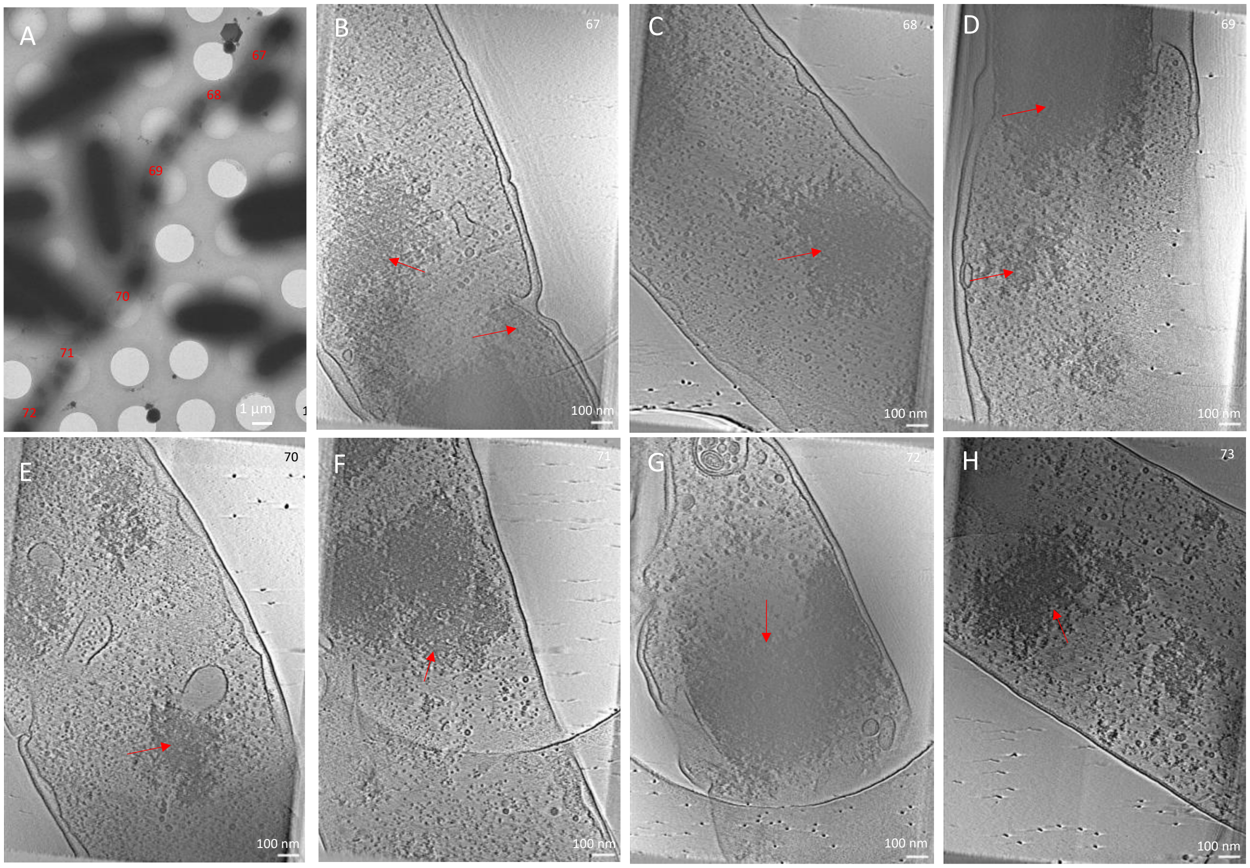


Figure S4. CET analysis of intracellular structures of *E. coli* strain expressing *pspA.* A filamentous *E. coli* cell of over 30 μm in length was used to take the tilt series (A) and reconstruct them into 3D tomogram (B to H). Aggregates are indicated by red arrows.

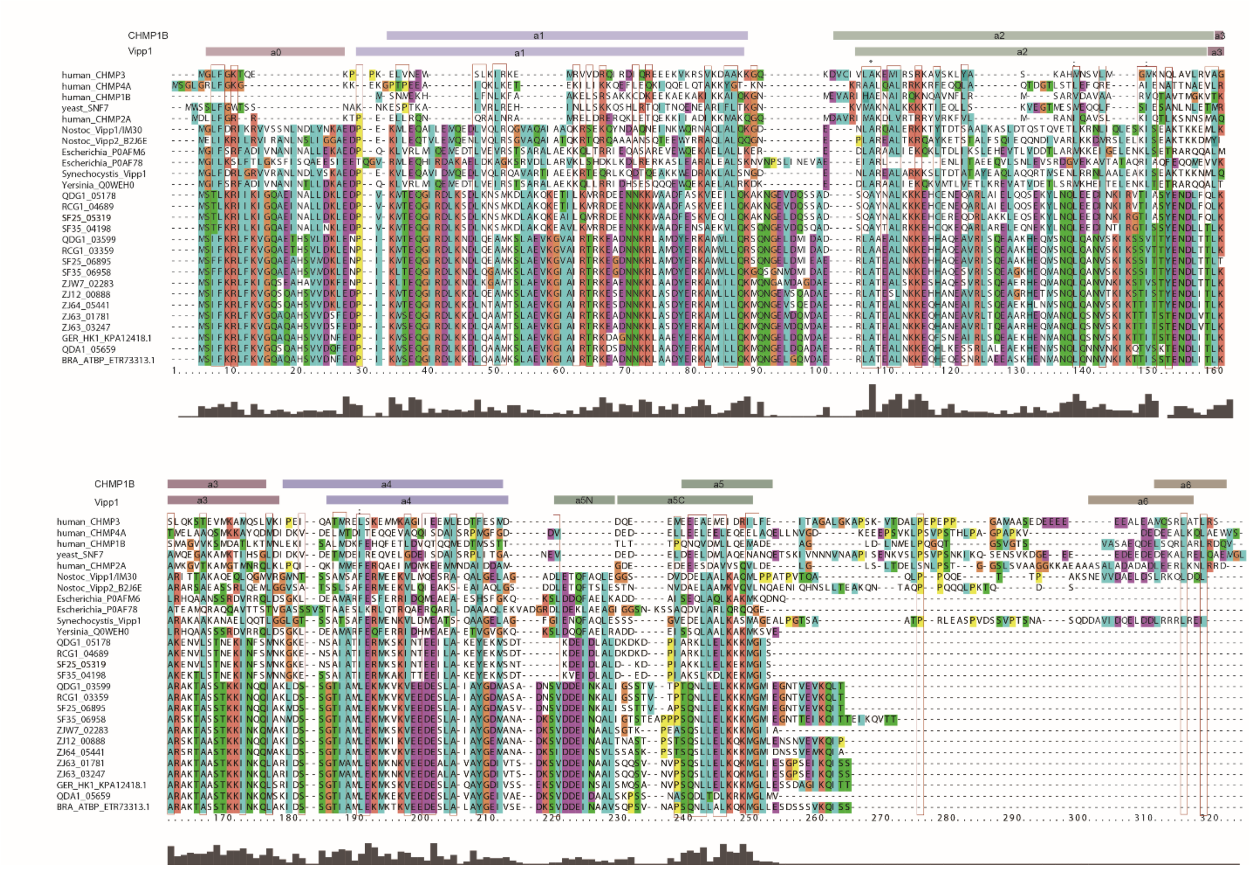


Figure S5. Multiple alignment of PspA proteins.

Sequences were aligned using Clustal Omega and include Vipp1/IM30 *Nostoc punctiforme* (Uniprot code B2J6D9), Vipp2 *Nostoc punctiforme* (Uniprot codeB2J6E0), *Escherichia coli* PspA (Uniprot code P0AFM6 and P0AFM6 ), *Synechocystis* sp. Vipp1 (Uniprot code A0A068MW27), *Yersinia pestis* PspA (Uniprot code Q0WEH0), human CHMP2A (Uniprot code O43633), yeast SNF7 (Uniprot code P39929), human CHMP1B (Uniprot code Q7LBR1), human CHMP4A (Uniprot code Q9BY43), human CHMP3 (Uniprot code Q0Y3E7) and pspA of MMPs. Cartoon shows Vipp1 and CHMP1B secondary structure.

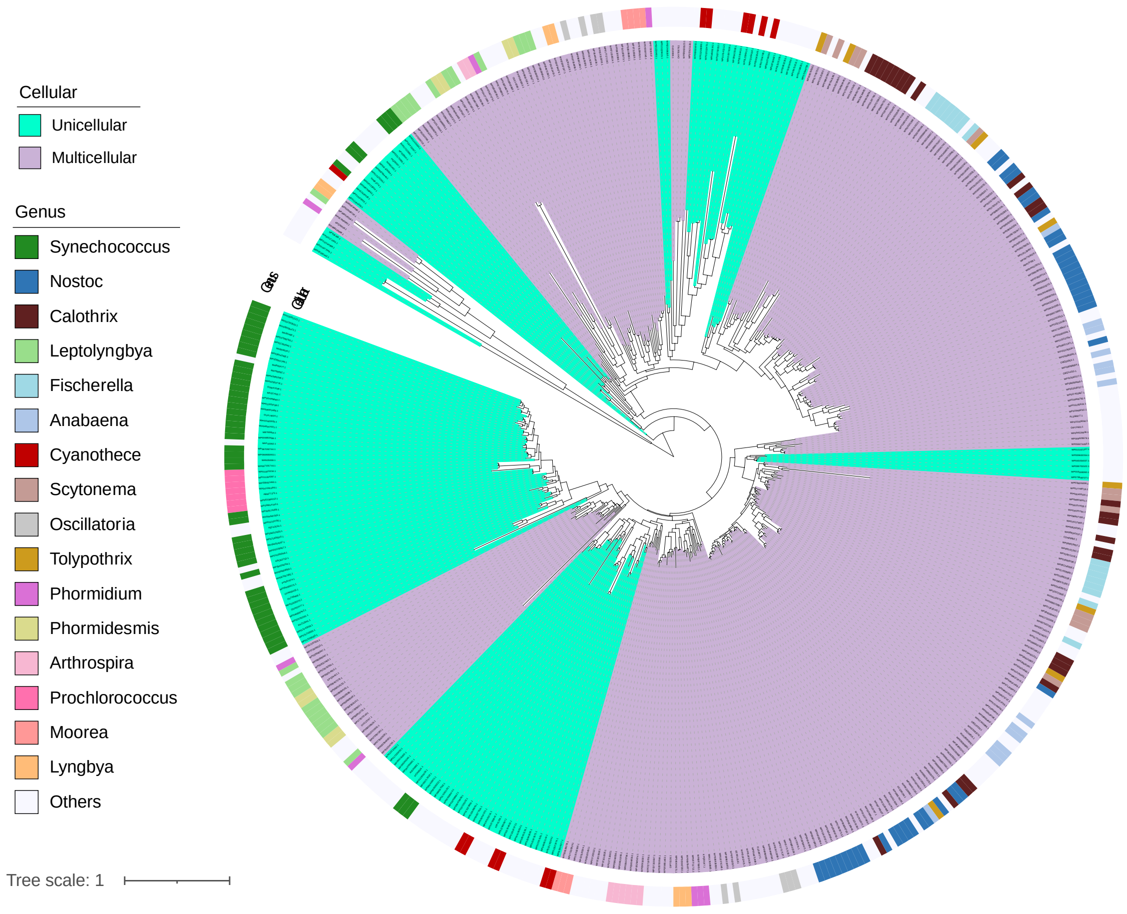


Figure S6. Phylogeny of cyanobacteria PspA.

Phylogenetic analysis of 298 *pspA* genes detected in 155 multicellular cyanobacteria genomes and 147 *pspA* genes detected in 114 unicellular cyanobacteria genomes. The inner clade color pale lilac, pale green mean *pspA* gene from multicellular or unicellular cyanobacteria, and the outside circle colored the genus of the *pspA* gene.

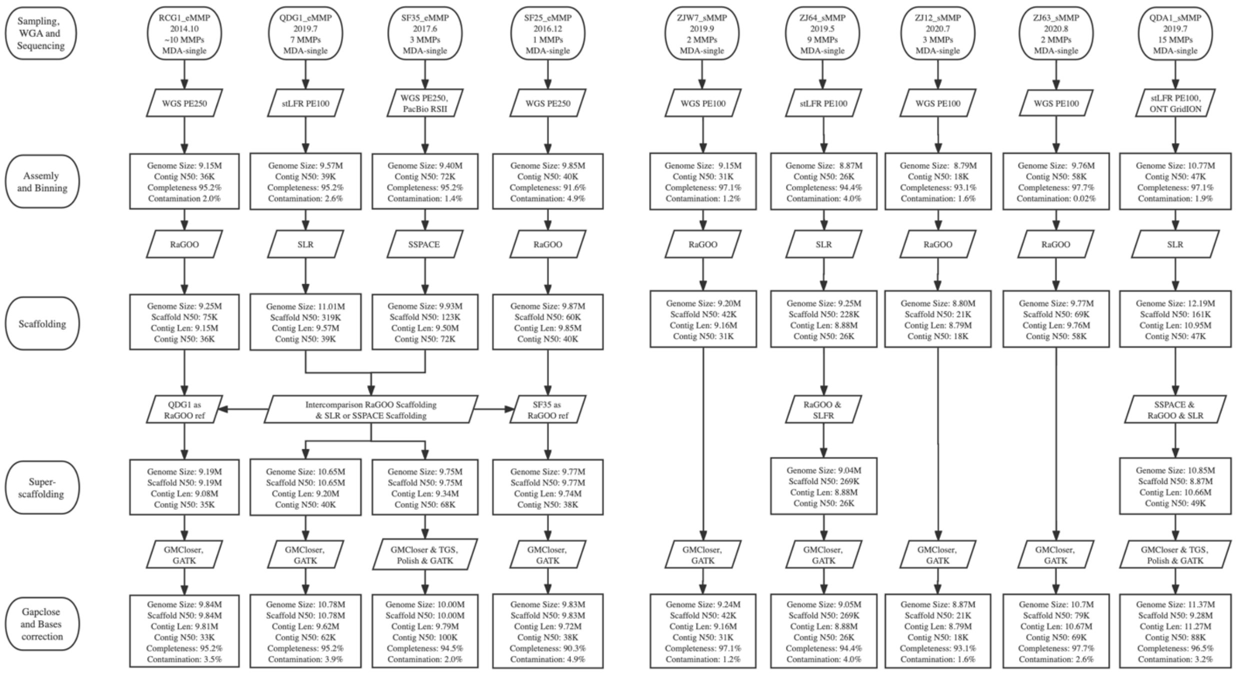


Figure S7. Sequencing, assembly and genome recovery flow chart of the study.

The left rounded rectangles showed the main processes of magnetoglobules samples, and different steps were connected by arrows. For details about the software, please refer to Materials and Methods. Figure created by ProcessOn.

Table S1. Pairwise comparison of 16S rRNA genes and average nucleotide identity (ANI).

| | RCG1 | QDG1 | SF25 | SF35 | ZJ64 | ZJ12 | ZJW7 | ZJ63 | QDA1 | GER_HK1 | BRA_ATBP |
|---|---|---|---|---|---|---|---|---|---|---|---|
| RCG1 | - | 100.00 | 99.16 | 98.45 | 91.17 | 91.19 | 90.60 | 92.39 | 91.76 | 92.99 | 91.56 |
| QDG1 | 99.59 | - | 99.16 | 98.45 | 91.17 | 91.19 | 90.60 | 92.39 | 91.76 | 92.99 | 91.56 |
| SF25 | 90.84 | 90.48 | - | 98.07 | 91.51 | 90.98 | 90.68 | 92.20 | 91.38 | 92.80 | 91.30 |
| SF35 | 83.07 | 83.01 | 84.33 | - | 91.38 | 90.74 | 90.60 | 92.20 | 91.32 | 92.54 | 91.51 |
| ZJ64 | 69.00 | 69.47 | 69.02 | 69.01 | - | 95.13 | 90.70 | 90.50 | 90.86 | 90.61 | 90.83 |
| ZJ12 | 69.34 | 69.46 | 69.27 | 68.93 | 76.58 | - | 90.78 | 90.77 | 91.19 | 90.86 | 90.64 |
| ZJW7 | 68.13 | 68.49 | 68.05 | 68.33 | 71.49 | 69.95 | - | 91.46 | 90.71 | 91.15 | 91.08 |
| ZJ63 | 68.62 | 68.90 | 68.47 | 68.32 | 70.13 | 70.07 | 70.98 | - | 96.20 | 96.93 | 96.41 |
| QDA1 | 68.85 | 70.06 | 68.70 | 68.80 | 71.35 | 70.49 | 73.91 | 73.69 | - | 96.59 | 98.07 |
| GER_HK1 | 69.18 | 69.75 | 69.14 | 69.11 | 71.52 | 71.29 | 72.11 | 74.45 | 74.81 | - | 96.32 |
| BRA_ATBP | 69.02 | 69.02 | 68.58 | 68.89 | 70.02 | 70.42 | 71.01 | 73.37 | 76.38 | 75.37 | - |

Table S1. Pairwise comparison of 16S rRNA genes and average nucleotide identity (ANI).

The four eMMP genomes are the ellipsoidal magnetoglobules from Rongcheng city (RCG1), Qingdao city (QDG1) and six-fours les plages (SF25 and SF35). The seven sMMP genomes include spherical magnetoglobules from Zhanjiang city (ZJ64, ZJ12, ZJW7 and ZJ63), Qingdao city (QDA1) and those reported by Kolinko et al. 2014 for GER_HK1 and Abreu et al. 2014 for BRA_ATBP_sMMP. The 16S rRNA gene identities and ANI comparison values were highlight with orange and blue color, respectively.

We propose the following taxonomic names using the cutoff values as previous described (Same species: 16S rRNA gene identity > 97%, ANI > 95%, RED > 0.85, Different genus: 16S rRNA gene identity < 92%, ANI < 83%, 0.7 < RED < 0.85).

Movie S1. PspA2 mediated formation of membrane sacculus.

Cryo-electron tomography of a *E. coli* cell expressing ellipsoidal magnetoglobule *pspA2* gene. Membrane sacculus can be observed at the cell pole.

Data S1. Metadata summary of analyzed genomes, PspA, MamB and MamM used in Figures 3, 4 and 5.

The CNSA or NCBI accession number of 57 genomes (included 11 multicellular magnetoglobules, 44 unicellular MTB and 2 related non-magnetotactic Desulfobacteraceae bacteria) used in Figure 3 were shown in column D. The PspA, MamB and MamM sequences were deposited into CNSA under the accession number CNP0003599. The sequence id of MamB, MamM genes used in Figure 4 were shown in column I and K, respectively, and PspA genes used in Figure 5 were shown in column G.
